# ALTAA: analysis of long-term activity patterns in ant colonies

**DOI:** 10.1101/2025.08.11.669637

**Authors:** Jinook Oh, Sylvia Cremer

## Abstract

Collective behaviours are a fascinating study area due to the emergent properties that can only arise in groups of interacting individuals. However, their quantitative study is often impaired by technical difficulties, creating either low-quality and sparse data, or impractical data amounts, particularly when capturing large groups over long periods of time. Common challenges arise from recording group members with as little obscuring of each other as possible, as well as in generating manageable data amounts with as high as possible information content. We here provide a multi-component system that allows to record, analyze and simulate the long-term spatiotemporal activity patterns of insect collectives, especially ant colonies. Our Ant Observing System, ALTAA, comprises a flat nest design to prevent occlusion of individuals, a recording system running on a low-power single-board-computer, and a set of computer programs performing quantitative analyses to guide the formation and validation of rules underlying the observed collective patterns. Our system is scalable in that it allows parallel, continuous observation of a high number of colonies using low memory space, with colony maintenance requirements (e.g. feeding, nest humidity) being achieved at lowest possible disturbance by the experimenter. We showcase the potential of the system in a study using the black garden ant, *Lasius niger*, where we analyze the spatiotemporal effects of different group size (1, 6, 10 ants), brood (larvae) presence or absence, as well as of different nest geometries, over a period of one week. We show that the ants’ motion activity has a weak periodicity in the range of 20 to 120 minutes promoted by larval presence, and that ants are spatially attracted to their larvae, the water source and the walls. We also find that the presence of nestmates lowers an individual ants’ motion activity. Observed data are compared to simulations of the temporal activity of the ants. ALTAA provides a powerful toolkit to quantify and interpret spatial and temporal collective activity patterns in (social) insects over extended periods.

## 1 Introduction

Social insects express a diversity of collective actions and behavioural changes in response to diverse stimuli they receive. In many cases, behaviours occur instantaneously as a clearly noticeable reaction to stimulus exposure, such as e.g. the immediate attack of an intruder to their nest [1, 2, 3], necrophoric behaviour to remove corpses [4], or sanitary care of pathogen-exposed colony members [5, 6, 7]. These clear behavioural reactions can be studied experimentally upon presenting the respective stimulus and observing the insects’ reaction. This approach has frequently been used to provide detailed observational data of a relatively low number of individuals or events, to describe the insects’ behavioural repertoire and its dependency on triggers. In addition, a whole-colony approach can be used to study how different stimuli change the social interaction networks of colonies [8, 9, 10, 11] or their spatial usage [12, 13]. Here, a common approach is to tag each individual group member before the experiment, by either RFID tags, colour coding or ARtags [8, 9, 10, 11, 14, 15, 16] and to follow the location of each individual over time. Individual tagging requires a lot of resources both during and after the experiment, yet it overcomes issues in social insect tracking – less prominent in less social species – in that their very close social interactions often lead to occlusion of individuals, introducing the constant need for differentiation of individuals during and after their social interaction. Social insects are thus ‘too social’ for the application of many conventional trackers developed for e.g. flies, fish, or mice [17, 18, 19, 20, 21], particularly when observed in experimental nest boxes with larger empty space than their natural nests, which are, in the case of ants, often tight underground tunnel and chamber systems [22]. We therefore implemented a tight nest space which allows the ants to move normally but constrains crawling on top of each other, to facilitate less resource-intense long-term observation of their activity without the need for individual tagging. Such methods are particularly valuable for research questions that do not require individual identity information but focus on the emergence of collective activity patterns in time and space. These can be analyzed by e.g. motion measures, as used in the analysis of collective phenomena [23, 24, 25]. It requires considerably less memory space so that even long-term observations over multiple weeks produce data quantities manageable for human analysis. Upscaling data size and managing it using combinations of non-linear computational units is needed to recognize complex patterns, however it often processes the data in a difficult way to be interpreted by humans. Both methods for investigating interactions among individuals and the global pattern of the group are required to understand a complex biological system. A further advantage of tag-free systems for long-term observations is that they are disturbance-free, are not affected by tag-loss issues and allow for birth and dead in the colony, hence allowing for studies of colony-development. Most importantly and other than the study of specific reactions to a particular trigger or of social interaction networks these analyses can also be performed in the absence of one-on-one interactions and the expression of particular behaviours and hence can capture a greater variety of activity patterns, including also more subtle changes within a noisy background. They hereby benefit from simultaneously extracting information of a high number of colony members (though they also can be used for the observation of individuals), over a long time-span and a potentially large nest area. Such subtle, gradual and hence seemingly chaotic behaviours that show large variances and form noisy patterns are difficult to analyze since they often occur as gradual changes of the activity level or of the intervals between intense activity bouts or of spatial usage patterns. Therefore, disentangling such patterns from the noisy background of stochastic behavioural activity in complex biological systems [26, 27] is not trivial, alike, for example, the study of magnetic waves or regional activations and neuronal connections in our brain [28, 29, 30]. In the social insects, such more subtle changes in activity or spatial usage may occur during the sensing or processing phase preceding a more specific behavioural action, such as e.g. perceiving a threat that will then lead to a concerted response. Moreover, just like most organisms, also insect colonies show a circadian rhythm [31, 32, 33], as well as shorter activity cycles [34, 35, 36], which require long-term analyses to be detected. Lastly, long-term analyses are also needed to determine the pace at which colonies go back from an alerted state (like e.g. after entering of an intruder [1]) to their baseline activity levels.

Therefore, despite the fact that many systems have already been developed to automate observations and recordings of insects using computer vision techniques [37, 38, 39, 40, 41, 42], many of which focus on individual tracking of tagged individuals, we felt that there is still need for a data generation and analysis toolkit for colony-level activity patterns that are inherently noisy and require integration of many data, including many replicates and long durations. We therefore developed ALTAA as a multicomponent system for automatized long-term observation, analysis and simulation of colony level activity in ants, with low computational effort and memory requirements. Our framework contains:

- AntOS: ant observation software and hardware to record the activity of individual or groups of ants,
- Visualizer: visualization software to plot and analyze the collected data, and
- AntSim: simulation software to model a theoretical rule-set to compare to the observed data.

## 2 Methods

### 2.1 Software & hardware tools for development

All the programs for this study were developed in Python (version 3.9, retrieved from https://www.python.org) with several packages; OpenCV (version 4.7, retrieved from https://opencv.org), wxPython (version 4.2, retrieved from https://wxpython.org), SciPy (version 1.1, retrieved from https://scipy.org) and Matplotlib (version 3.7, retrieved from https://matplotlib.org). Arenas for the observation/experiment sessions were designed in Blender (3.0, retrieved from https://www.blender.org) and printed using a stereolithography (SLA) 3D printer (Elegoo Mars 2, ELEGOO, China). The frame for housing the printed arena was made using aluminum beams named as MakerBeam (MakerBeam B.V., Netherlands). A developed software for observation/experimentation were installed and run on a single-board computer (SBC hereafter; Raspberry Pi 4b 4GB RAM, Raspberry Pi Foundation, United Kingdom). A camera (Arducam with Sony IMX477 sensor, Arducam, China) was attached to the SBC for the image acquisition.

The framework described in this study has three major components: (i) the recording of the behavioural observations during the experiment, accomplished by ‘AntOS’, (ii) the data analysis conducted by ‘Visualizer’ and (iii) a possible simulation using ‘AntSim’, with the latter allowing to run a series of simulations to infer possible rules underlying the observed data (see Figure 1).

**Figure 1.**
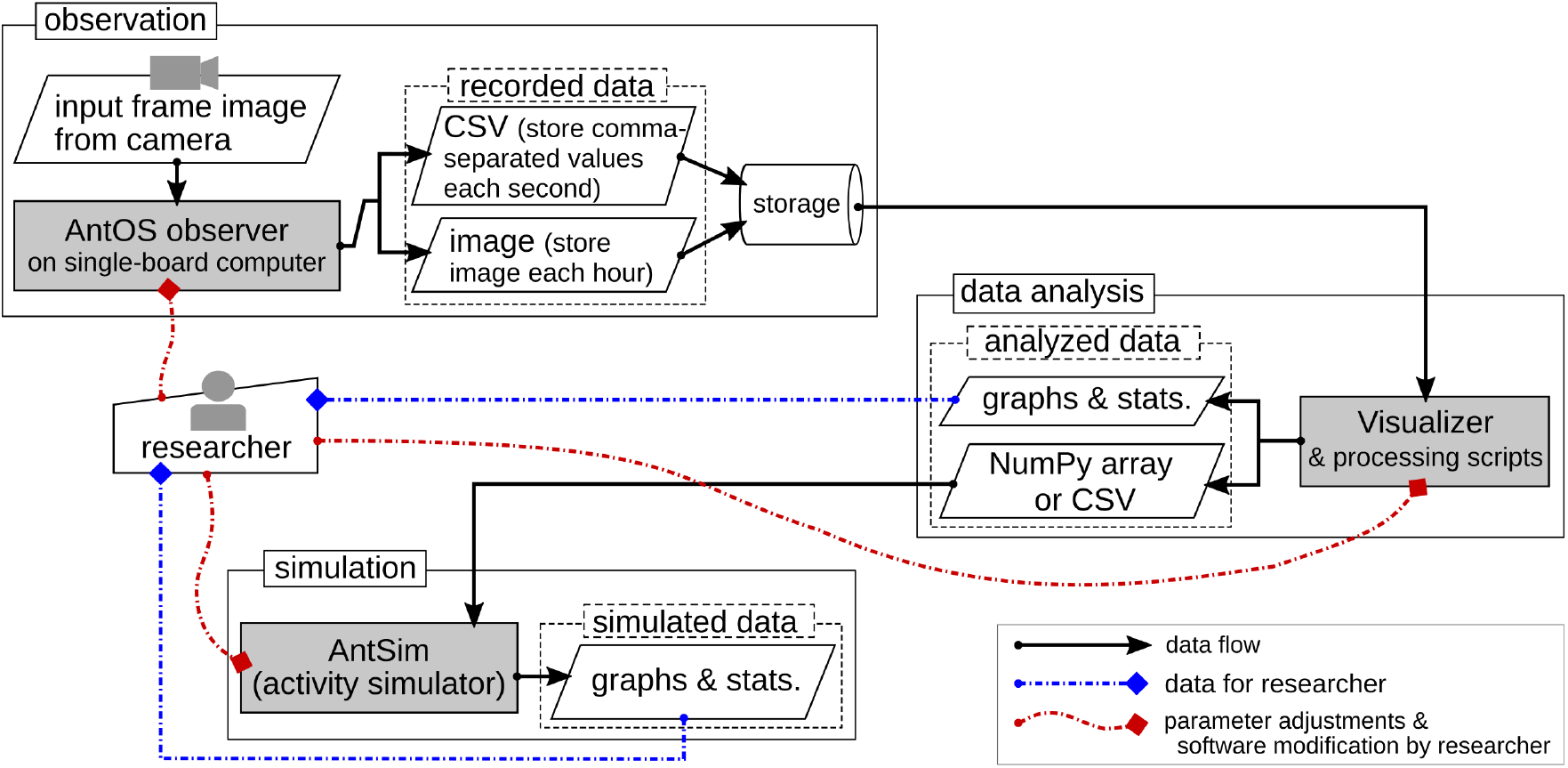
Concept diagram of the framework in this work

Current Python files comprising the aforementioned three programs, which belong to the group of code named ‘CremerGroupApp’, are depicted in Figure A1 in the Appendix. Some utility functions and classes are used for multiple programs. Functionalities of two files, ‘modFFC’ and ‘modCV’, are used for all programs, ‘modRasp’ is used for AntOS, and ‘graphFuncs.py’ is used for Visualizer and AntSim (see Appendix ‘Commonly used classes & functions’).

### 2.2 AntOS; Apparatus

We protected our experiments from environmental influences by them being performed within a box, built from a frame (MakerBeam) and six wooden panels. The outside of the wooden panels were painted with black matt paint to block outside lighting, while the inside was a bright ivory colour for indirect lighting. To further ensure homogeneous lighting inside the box, the light bulb inside was covered by a light-softening diffuser fabric. The light bulb was a non-flickering LED (12V, 1.2W, 120 lumen, 6000K), which was located at the corner of the apparatus avoiding a direct lighting to the camera lens or the Plexiglas lid of the flat-nest. It also contains a SBC for running AntOS software and acquiring data from a camera (via Camera Serial Interface) and a DS18B20 digital temperature sensor (via General Purpose Input/Output;GPIO pin on SBC); see Figure 2. We designed a ‘flat-nest’ (Figure 3) providing an arena for the observation, (i) to improve data accuracy by restraining occlusion of individual workers being on top of each other, (ii) to decrease the maintenance and analysis efforts, and (iii) to be able to create an arena with complex design. The height of the flat-nest in this study was designed for observation of the black garden ant, *Lasius niger* [43], considering the worker’s body size, approximately 1*mm* wide and 2 *–* 3*mm* long and a queen body size, approximately 2 *–*3*mm* wide and 9*mm* long.

**Figure 2.**
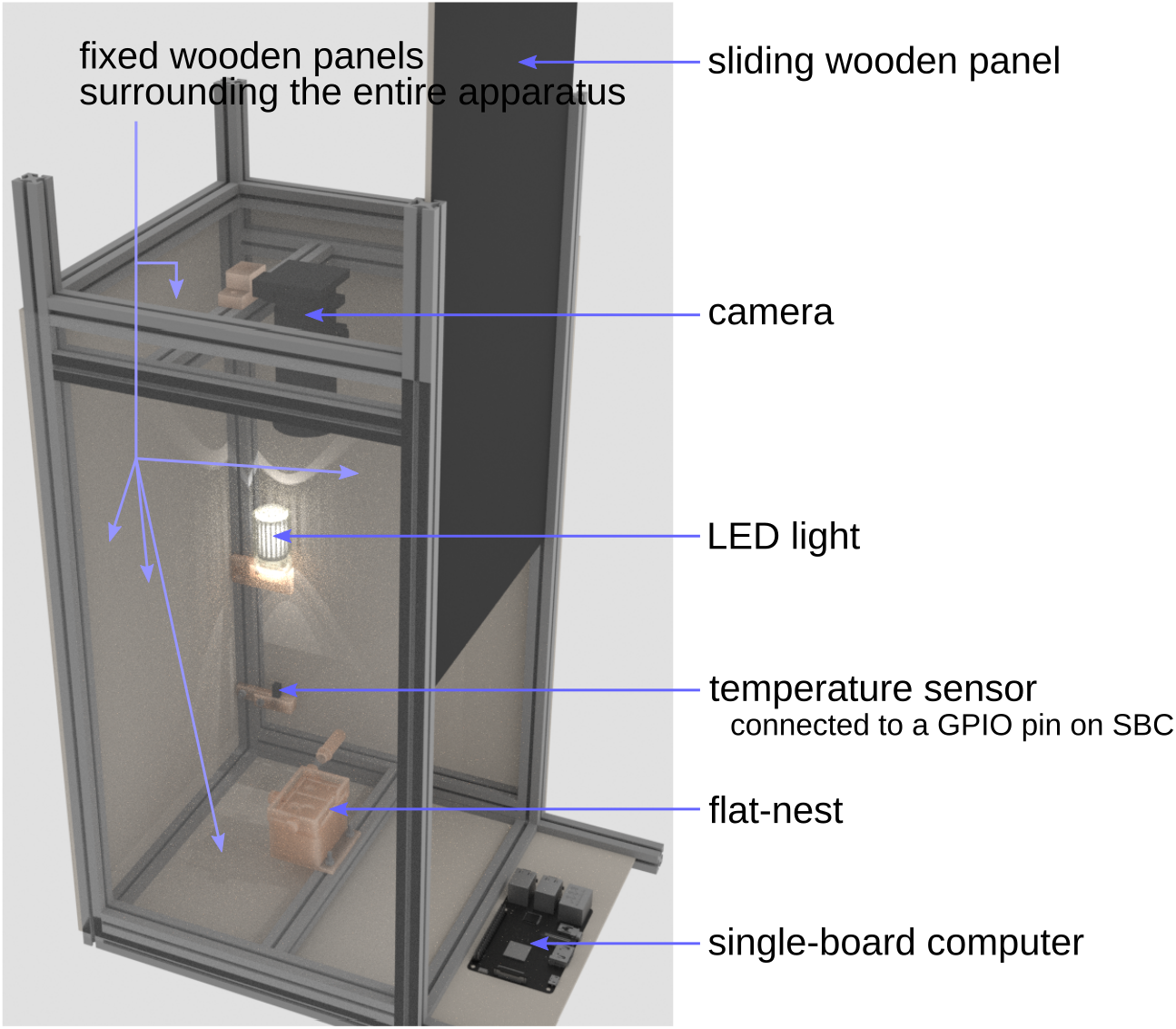
AntOS apparatus * Some of the wooden panels are rendered transparent to display components inside.

**Figure 3.**
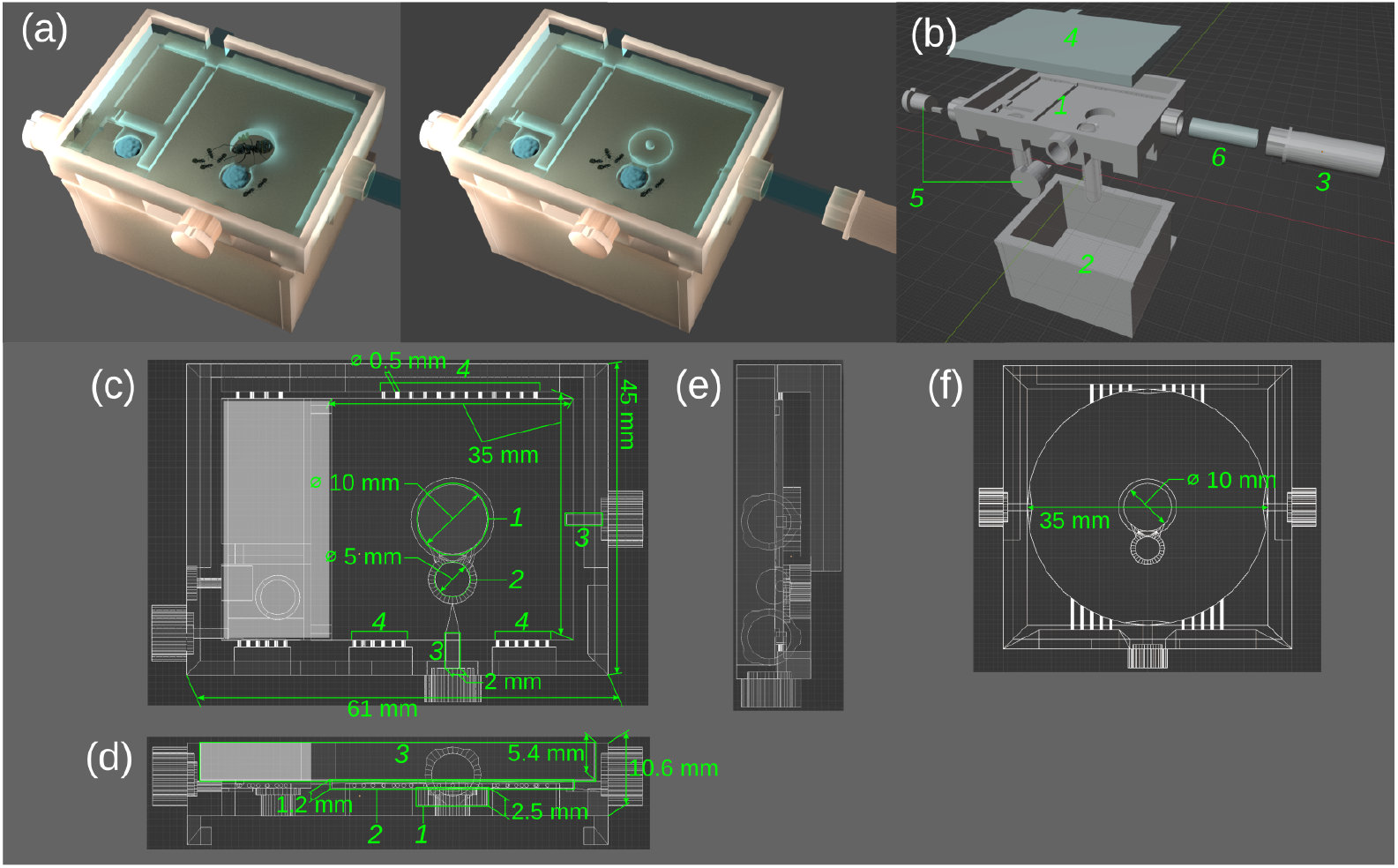
Flat-nest; (a) Overview of the flat-nest with and without a queen ant, (b) Major parts; *1*: arena, *2*: water container, *3*: sucrose solution container, *4*: transparent Plexiglas lid, *5*: plugs for outward passages, ø6: transparent rubber tube (6 mm), (c) Top view of a square-shape arena. A secondary chamber (greyed area on the left-side) was not used in the current study; *1*: circular indentation when housing a queen ant, *2*: hole for cotton string to provide water, *3*: outward passages, *4*: ventilation holes, (d) Front view of the arena; *1*: deeper area for a queen, *2*: shallow area for workers, *3*: place for transparent lid, (e) Side view of the arena, (f) Top view of the second flat-nest with circular-shape arena, used in the current study.

After testing multiple heights, we used 1.2*mm* of the internal space (Figure 3-(d)) of the arena, and an indentation of additional 2.5*mm* (leading to a final height of 3.7 mm) for the queen chamber. This setup restricted the queen to a central chamber, but allowed normal worker activity throughout the nest, yet restraining (but not making it impossible) workers moving on top of one another. At a higher height of 1.5*mm*, crawling over each other still occurred regularly, whilst a lower height of only 1*mm* restricted worker activity to certain degree.

The flat-nest can house the ants for observation during a long period such as weeks or months, due to its water container, sucrose container and ventilation holes (Figure 3-(b)-*2 & 3*, and Figure 3-(c)-*5*). The water is provided to the ants through a cotton string in a hole (Figure 3-(c)-*2*)(Figure 3-(c)-2), absorbing water from the water container (Figure 3-(b)-*2*). The ventilation holes prevent water condensation on the transparent ceiling, which could otherwise interfere with the image processing. For maintenance of the water and sucrose in the containers (in our case performed on a weekly basis), the AntOS program is paused while the containers for the sucrose solution and water to be detached and refilled.

Since the flat-nest is produced by 3D printing from resin, the nests have to be carefully post-processed after printing before use to ensure ant health. The toxic effect of the resin material in SLA 3D printing was already reported for zebrafish embryos [44], which could be mitigated by the post-processing using ultraviolet (UV) light.

To prevent toxic effects for the ants, which can show as reduced movement or activity, cessation of queen egg laying, and even death, the newly printed flat-nest should be (1) cleaned using isopropyl alcohol (IPA), (2) exposed to UV light, (3) water-washed and dried in a well-ventilated space, and (4) tested with a small number of workers. If any abnormal behaviour is noticed, the above steps should be repeated. The length of the IPA cleaning and UV light exposure varies depending on the resin type and its production company. The FDM (Fused Deposition Modeling; filament type) 3D printing could be used with less safety steps, yet it has a reduced printing resolution compared to the SLA.

Moving a colony or multiple workers of *L. niger* into the flat-nest could be difficult due to the low height of the arena. We propose to first add the queen and brood into the open nest, and then to move workers into the closed arena using a funnel and a tube (see Figure 4). For this procedure, a transparent rubber tube is prepared with one side blocked with a cotton ball and another side with a funnel attached, which is also 3D-printed with the matching diameter to the tube. To prevent the ants from climbing out of the funnel, Polytetrafluorethylen (PTFE, like Fluon) can be applied to its interior walls. To first trap the ants in the tube, they can be dropped into the funnel using tweezers, then the funnel is tapped to make them fall into the tube. Then, the upper end of the tube is blocked by another cotton ball. After removal of the funnel and the first cotton ball, the tube is quickly connected to the arena. The ants in the tube are encouraged to move into the arena by softly pushing the remaining cotton ball. We found workers to typically quickly move to and stay in the arena, also being attracted by the wider space and the brood items in the arena. After ant integration, the passage can be either blocked with a printed plug, or connected to another section such as a sucrose container or a chamber.

**Figure 4.**
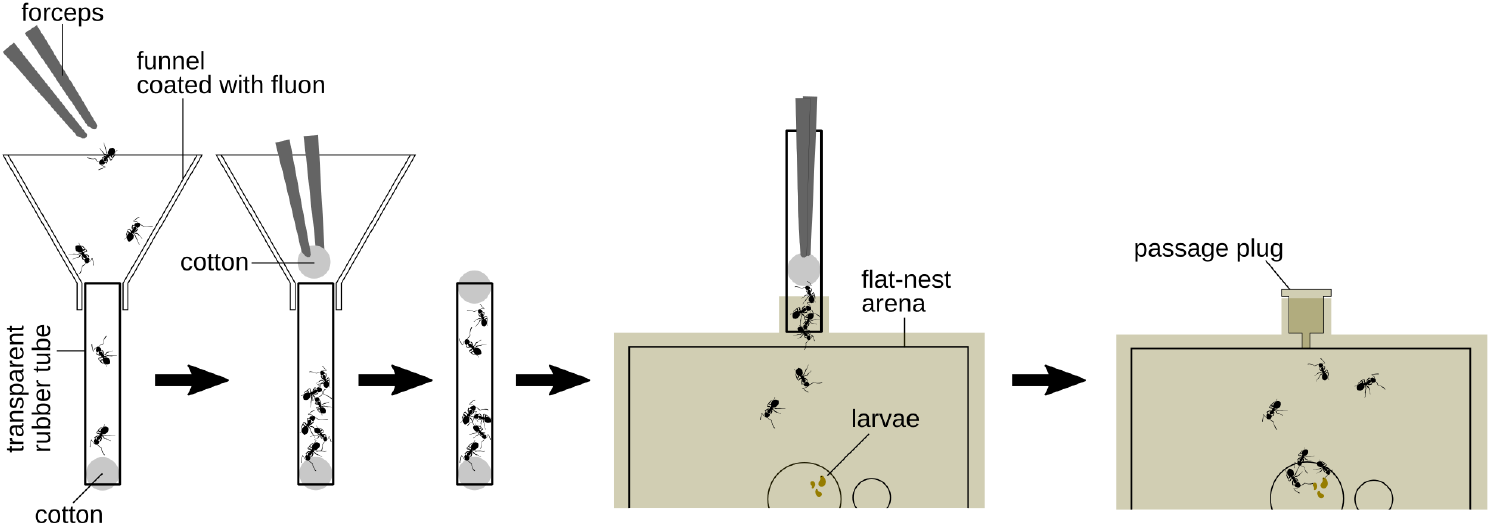
Introducing ant workers into the flat-nest arena

### 2.3 AntOS; Software

The AntOS program collects the data from the camera and other sensors in apparatus, like the temperature sensor. In the graphical-user-interface (GUI) of AntOS, the user can adjust several options including the camera index used, the session duration, the regions of interest (ROI), intervals for the temperature logging and saving ROI images to files, colour values in HSV colour space (Hue, Saturation and Value) for detecting the inherent ant colour or an applied colour marker. The number of ROIs, which can be set in the frame image, depends on the size of the flat-nest and the camera type. In the current study, we set ROIs to three. Upon starting it, the program (i) reads and processes a frame image from the camera every second to save a resultant data in a CSV data file for each ROI, (ii) records the temperature inside of the AntOS apparatus every 10 minutes (adjustable) to a log file, and (iii) saves an image of each ROI as a JPEG image file every hour (adjustable). The image files and the resultant CSV data files for each ROI are saved in a separate folder.

Frame processing is used to detect motion to store center points of motion segments, detect ant colours to store its rect (rectangle denoted by x, y, width and height) info and a possible additional colour code applied to the ants (not performed in the current study) to store its position. For the motion detection, AntOS calculates the difference in consecutive frame images, finds contours from the difference between images, calculates the center point of each contour and saves the x/y coordinates of contours with a timestamp in the resultant CSV file. For the ants’ colour detection, it detects blobs with the user-defined HSV colour and stores the rect of each blob’s contour, yet AntOS can be used without this additional information only based on motion data, as in the experimental study described below.

The temporal resolution of the frame processing is set to one second due to the time needed for frame retrieval of the SBC from the camera and running the processing Python code. AntOS on the currently used SBC takes 0.5-0.6 seconds to retrieve the frame image (1920 × 1440 resolution) and an additional short time (depending on the number of blobs that moved in each frame) to run the image processing code. The part (ROI) of the frame image is saved as an image file every hour to allow the experimenter to check for possible artifacts or disturbances to the system, such as a software error or physical disturbance of the apparatus during the session, which would affect the resultant data. The filename of each saved image is a timestamp. By following use of the Visualizer program (described below), motion center-points, occurred at certain time, can be drawn on the ROI image that was saved at that exact time. It draws all the corresponding motion points to all the saved images at each hour so that the user can check each motion point is drawn on the ants in the images.

The AntOS code is split into three files, considering the code modularity, ‘antOS.py’ for the GUI, ‘modSessionMngr.py’ for calling functions on the scheduled time during the session and ‘modRasp.py’ to retrieve the frame image and send/receive data to/from sensors/actuators via GPIO pins on the SBC. For different functionalities depending on the experimental setup, the image processing can be modified in ‘modSessionMngr.py’, and managing sensors and actuators attached to GPIO pins can be modified in ‘modRasp.py’.

### 2.4 Data exploration/visualization; Visualizer

The Visualizer program is to draw graphs and images (such as heatmap images, or screenshots) from the data obtained by AntOS. These observed data from the experiment can later be compared to theoretical rules of ant behaviour by producing processed datasets in a NumPy [45] array for the simulation software AntSim, detailed below. The main functions of the Visualizer program lie in the analysis of motion-point data, as showcased in our experimental study below. After selecting a data folder, the user can select a graph/image type to draw, set different parameters and press the ‘draw’ button in the GUI to start bundling data with a user-defined interval (such as several seconds or minutes to be represented as a single data point in the resultant graph) and drawing a graph with the bundled data. It offers the following options to draw different graphs or images:

- *initImg* shows the first image of all saved ROI images. This is for setting parameters directly from the image. On the loaded image, the user can mouse-click/drag to select certain points such as a brood/food/garbage position, the region of interest, and an area of grid-cells.
- *sanityChk* is for the user to check if motion points occur only on the ant’s body with the saved ROI images to detect possible artifacts.
- *intensity* draws a graph with the number of motions in each bundled data point.
- *intensityL* draws a graph of the number of motions around three user-defined locations; brood, food and garbage.
- *dist2b* draws a graph with measurements of the maximum (among all motion points in the data frame) distance between the brood position and a motion point in the bundled data.
- *intensityPSD* draws the power spectral density graph using Welch’s method [46] on the data processed with ‘*intensity*’.
- *spAHeatmap* draws a heatmap image with the motion points.
- *spAGridHeatMax & spAGridHeatMean* draws a graph with the max/mean value in each grid-cell of the heatmap during a given time (bundle-interval).
- *proxMCluster* draws a graph with the motion cluster value, which represents multiple ants moving next to each other for a certain duration (*≥* 3*sec*.). It is described with a pseudo-code in Algorithm 1 in Appendix.

### 2.5 Simulation; AntSim

With AntSim, the user can implement theoretical rules of ant behaviour, based on hypotheses of ant decision rules or inferred from the processing of the observed experiment in Visualizer. AntSim simulates motion data in the same temporal resolution and format as the raw real-world ant data acquired by AntOS. The resultant graphs from the simulated data can be directly compared to the output of Visualizer using the real-ant data to evaluate the validity of the theoretical underlying rules for ant behaviour. The main focus of the current AntSim is to simulate temporal activity patterns of ant groups, following the simulation procedure outlined in Figure 5 with a set of user-defined parameters: length of data, data-bundle interval, interaction of workers and threshold ranges to boost the distribution of the activity & inactivity durations.

**Figure 5.**
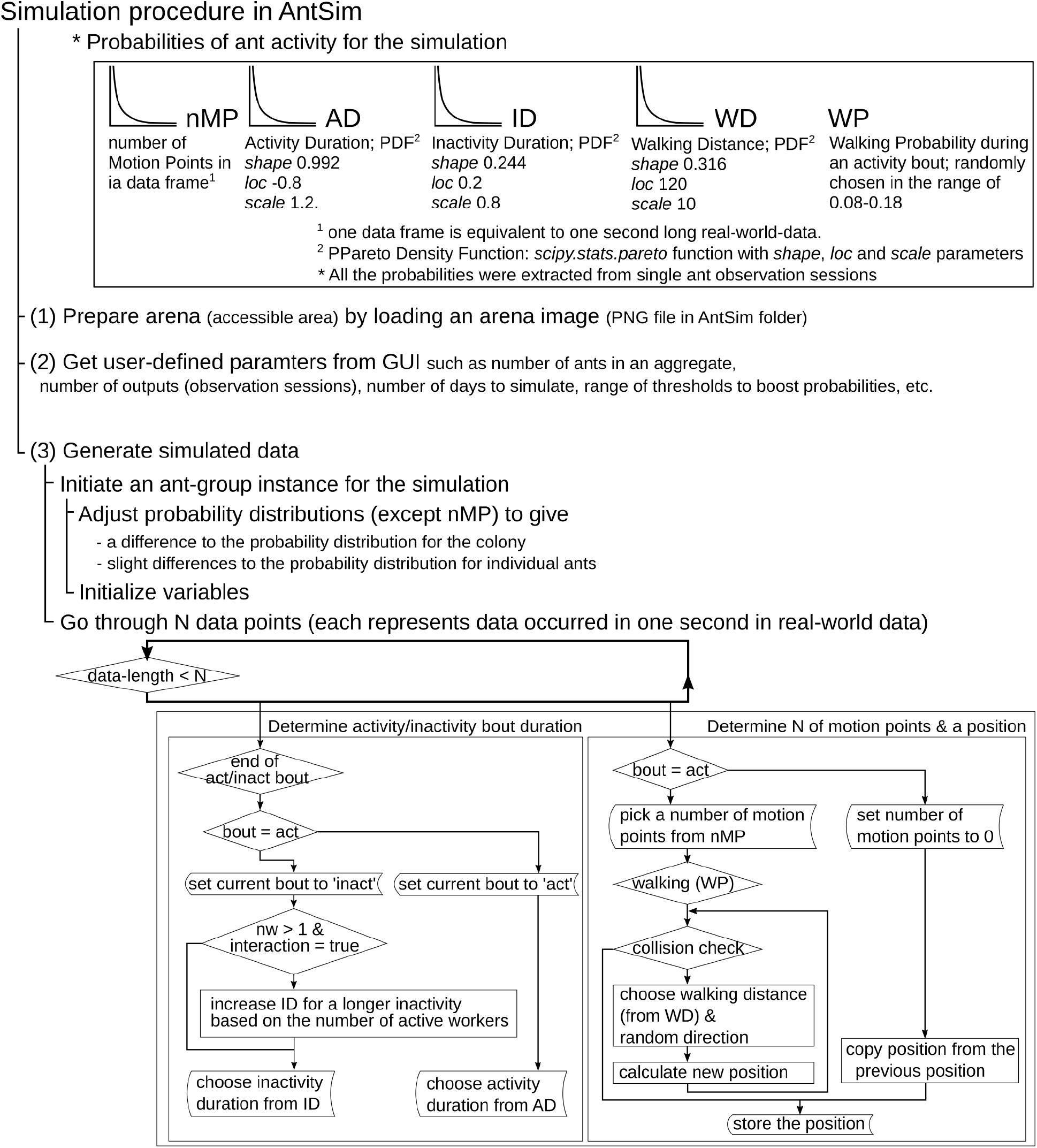
Simulation procedure in AntSim

We have implemented two major simulation types (i) ‘*n[N]_[i or s]]*; where *N* is the number of ants, *i* is for interaction among ants and *s* is for non-interactive ants’, and (ii) ‘*mComp_[i or s]*’ When the simulation procedure is finished, the program outputs an intensity barplot, its power spectral density, boxplots of the activity and inactivity durations, and heatmap images. In ‘*mComp_[i or s]]*’, ant groups with the same parameters, but of different group size (one, six & ten), are simulated. With this simulation option, intensity barplots, power spectral density graphs, boxplots of the activity and inactivity durations, and heatmap images are produced for each group size. Additionally, a comparison graph for the number of movements by group size is produced.

### 2.6 Experimental study

We were here interested in analyzing the spatial and temporal activity patterns of *L. niger* workers when alone or in a group of six and ten ants, in the presence or absence of brood and in different nest architectures. To this end, eleven colonies of this monogynous and monomorphic species common to the Northern hemisphere, were reared from queens collected during a large nuptial flight (which regularly occur from July to August [43, 47]) in Klosterneuburg, Austria in July 2022. Collection of this unprotected ant species was performed in compliance with international regulations, in particular the Convention on Biological Diversity and the Nagoya Protocol on Access and Benefit Sharing (ABS), and all experimental work followed European and Austrian law and institutional ethical guidelines. After collection from the field, the queens were reared in the dark to a fully grown colony with workers in plastic boxes (Tupperware) containing a constant supply of water, 1*M* sucrose solution and dried mealworm powder. Routine maintenance work, including providing water and food and cleaning, was performed every week.

We analyzed two experimental setups:

1) To test for the effect of group size on activity, one, six and ten worker ants per colony were sampled randomly (except for avoiding freshly emerged callow ants) and housed in one flat-nest each of the square arena type (n1, n6 and n10; Figure 3-(c)). Several mid to late stage larvae from the same colony had been added per replicate. The motions of the three group sizes per colony (n=11) were recorded for one week each between September and November in 2022.

2) To test for the effect of brood presence and nest architecture, we repeated the experiment one year later (from July to October in 2023) for the sizes of one and six workers (n1* and n6*), to compare these new setups without larvae and being placed in a round instead of square arena to the previous n1 and n6 replicates (see Figure 3-(f)).

We collected data by AntOS, analyzed it by Visualizer and simulated temporal activity by AntSim. Statistical comparisons (Wilcoxon Signeed Rank and Rank Sum Tests, Kruskal Wallis Tests) were performed in R (version 4.1.2). In pairwise posthoc comparisons and due to reuse of the n1 and n6 data with larva for their comparison to the n1* and n6* larvae-free situation, Bonferroni corrections were used to adjust for multiple testing. Two-sided, adjusted p-values are reported.

## 3 Results

### 3.1 Intensity; number of motion points

We first analyzed activity ‘intensity’ denoted by the number of motion points in a given duration, with example graphs being shown in Figure 6-(a). When comparing the intensities between the group sizes of six vs ten ants, the n10 motion was lower than expected by sheer upscaling from the six ants group size (*m/*6• 10, where *m* was the number of motions of n6 from the same colony; median 0.896 instead of the expectation of 1.0; Wilcoxon rank sum exact test, two-tailed test, W = 22, p = 0.01). Therefore, an increase by four additional workers led to a reduced motion. Note that we derived the expected value for the 10 ants only from the six ants and not from single ants, as the latter may show substantial individual differences.

**Figure 6.**
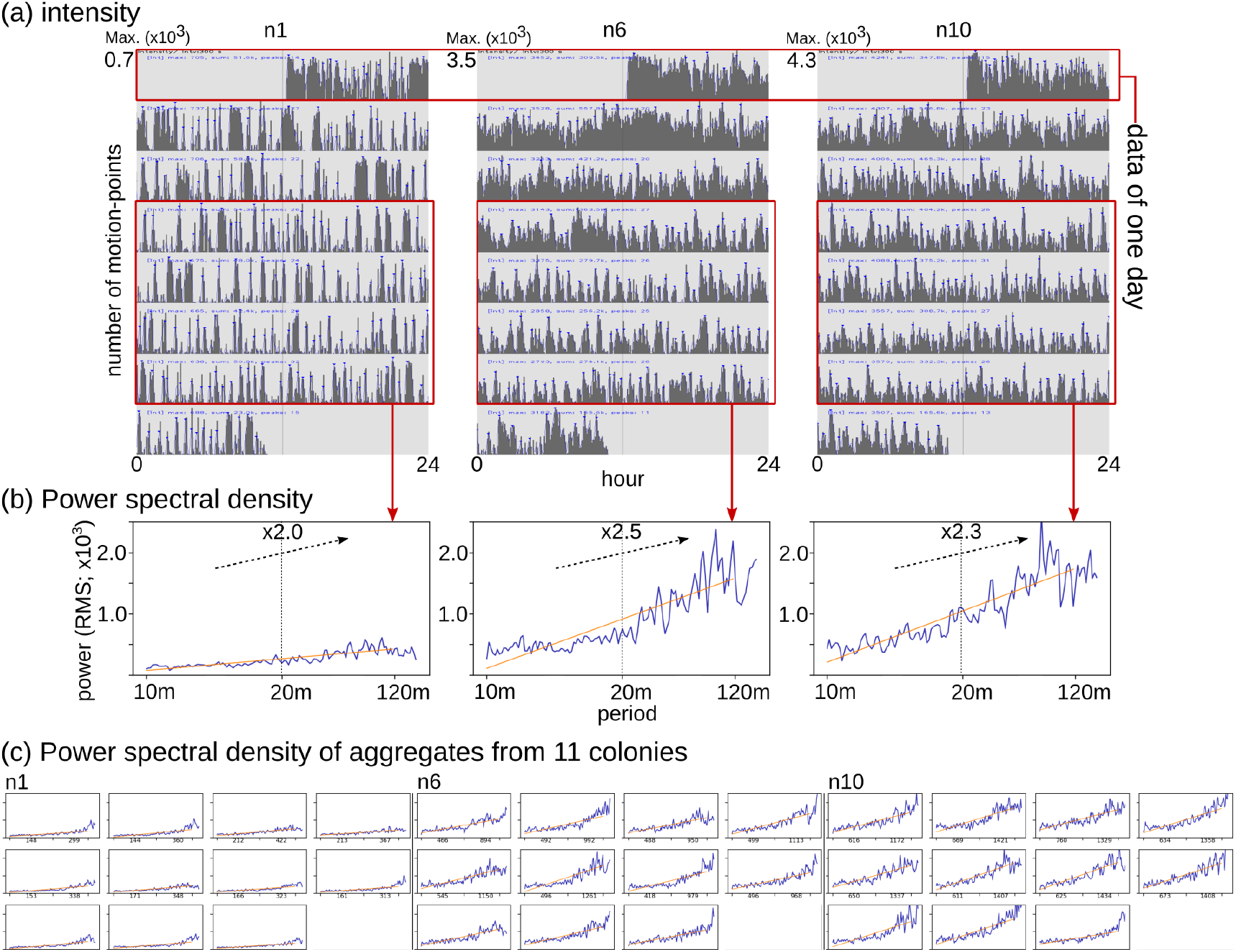
(a) Activity intensity (number of motion-points); n1, n6 & n10, as exemplified from one colony, (b) Power spectral density graphs of the intensity data shown in a (from four days in the middle of the session). The values on the y-axis are root-mean-square values and the float number in the middle of each graph indicates the fraction of mean power values of longer to shorter period. (c) Power spectral densities of n1, n6 & n10 for all 11 colonies

When comparing same-sized groups with and without larvae being present, we could not detect any significant differences in the sum of intensity (Wilcoxon signed rank exact test; n1 vs n1*: V = 51, p = 0.123, n6 vs n6*: V = 48, p = 0.206).

### 3.2 Temporal pattern

To describe the temporal activity pattern or periodicity of the ants, we calculated the power spectral density for the different group sizes (n1, n6 and n10) per colony. In contrast to *Leptothorax* ants (*L. allardycei* [34] and *L. acervorum* [48, 36]) that show activity peaks every 15-30 minutes, we found no synchronized short-term activity peak in *L. niger* ants in our experiment. Yet, many unconcerted peaks of activity occurred within the period of 20-120 minutes. The mean power in this long duration range of 20-120 min. was approx. double than that in the short duration range of 10-20 min. (Figure 6).

The resulting fractions of *∼* 2.0 were consistent across group sizes (Kruskal-Wallis rank sum test; chi-squared = 0.076, df = 2, p = 0.963). However, the fraction dropped when there were no larvae, reflecting that the presence of larvae stimulates the worker ants’ activity to be more periodic in the long duration of 20-120 min (Figure 7, Wilcoxon signed rank exact test; n1 vs n1*: V = 66, p < 0.001, n6 vs n6*: V = 60, p = 0.014). Note that this fraction drop occurred despite the fact that overall activity intensities did not differ in the presence or absence of larvae (see section 3.1).

**Figure 7.**
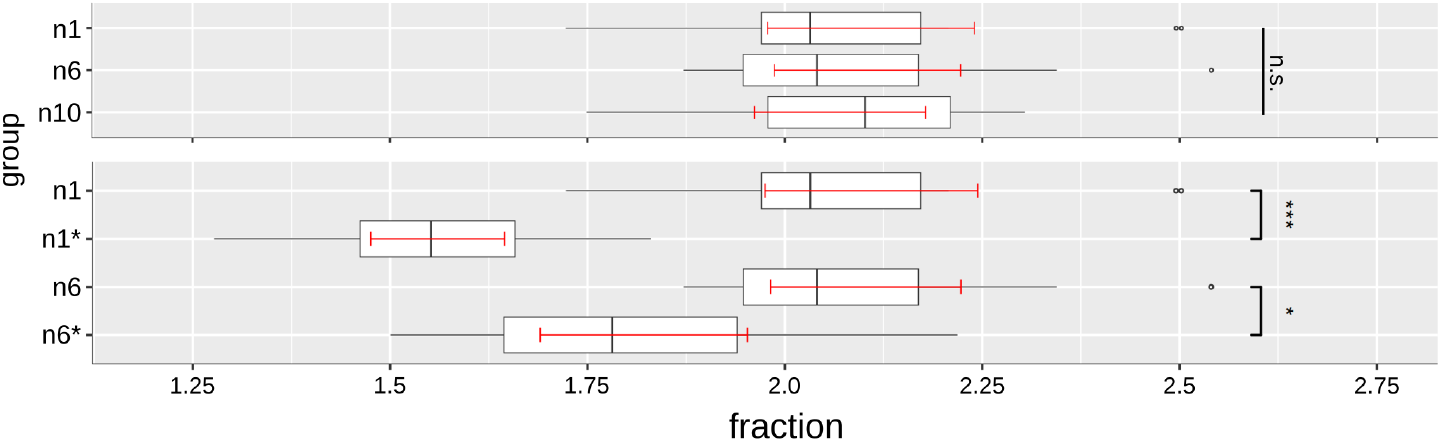
Fractions of mean power in the long duration range over the short duration range (a) for the three group sizes with larvae (n1, n6 and n10), and (b) for same-sized groups with and without larvae (n1 vs n1*, n6 vs n6*). Boxplots show the median and interquartile range (IQR). Whiskers extend to the most extreme data points within 1.5*× IQR* of the quartiles; points beyond this range are plotted as outliers (dots). Red error bars indicate the 95% bootstrap confidence interval of the mean. Significant differences between contrasts are depicted as * for p<0.05, *** for p<0.001, and n.s. for non-significant. Data from all 11 colonies.

### 3.3 Spatial pattern

To analyze the ants’ spatial usage preferences, we used Visualizer to create heatmaps and graphs (Figure 8). By comparing same-sized groups with and without larvae respectively, we found that workers are strongly attracted to larvae by showing higher activity in the center area in the presence of larvae (Figure 8-(a) and (b)), and performing a more localized activity with larvae (Figure 8-(c)). Figure 8-(b) shows that the attraction point of groups with larvae (n1, n6 and n10) were all quite close to the center, while groups without larvae (n1* and n6*) rather stayed away from the center.

**Figure 8.**
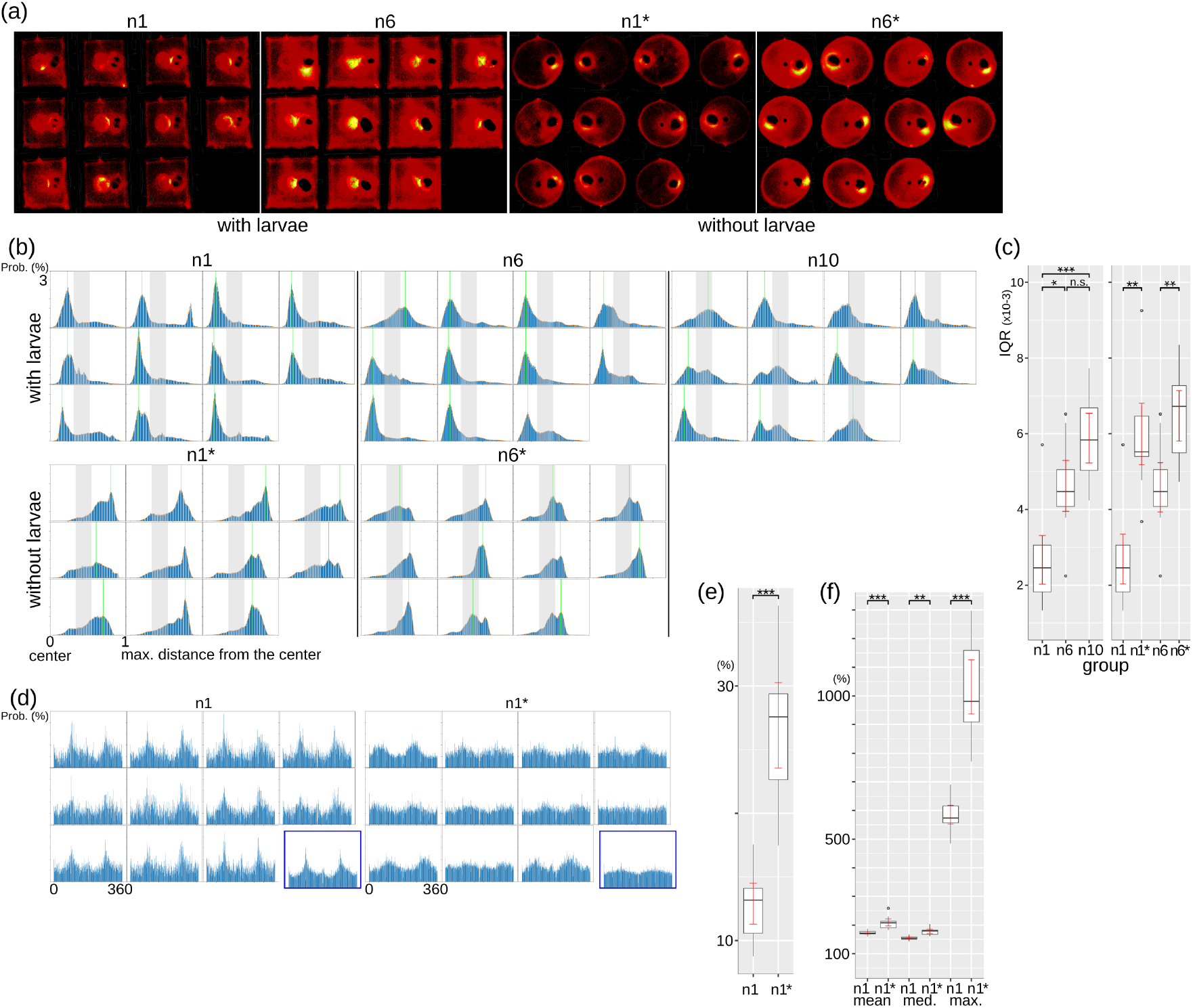
Patterns of ant spatial usage shown as (a) heatmap of motion-points in single ants or groups of 6 ants, both with and without larvae and in the square vs circular arena (yellow indicates higher activity than red, black indicates no activity), (b) distributions of the distance (from the arena center) of each colony in n1, n6, n10, n1* and n6* (grey zone depicts the distance to the water cotton ball and the green line depicts the highest peak distance), (c) interquartile range of analysis (b), with single ants and groups or 6 compared both by size and presence/absence of larvae, (d) direction probability of single ants when walking in the square vs circular arena (distance of two motion-points in two consecutive frames exceeding 120% of the ant’s body length). The last graph marked with blue rectangle for each group size displays the mean probability distribution, (e) the percentage of walking in single ants, and (f) a comparison of the single ants’ walking distance (in terms of ant’s body length) with and without larvae. Boxplots are formatted as in Figure 7.

Also, the ants are attracted to edges or wall-like structures, as groups of all sizes show high activity around the edges of the arena (Figure 8-(a)). Figure 8-(d) shows that ants walk along the edges, i.e. they have a higher probability of walking to the perpendicular directions in the square arena compared to the circular arena (Figure 8-(d)). Moreover, they seem to be attracted to the water-source (high activity of n1* and n6* around the water source (Figure 8-(a)).

The interquartile ranges (IQR) of the probability distributions of n1, n6 and n10 (Figure 8-(c)) reveal that activities are more dispersed in groups of 6 or 10 ants compared to single ants (Kruskal-Wallis rank sum test; chi-squared = 18.131, df = 2, p < 0.001; pairwise Dunn’s test for posthoc comparisons with Bonferroni correction: n1 vs n6, p = 0.036, n1 vs n10 p <0.001, whilst n6 vs n10, p = 0.256). Comparison of the IQRs of single ants resp. groups of 6 ants with and without larvae, revealed that the ants showed more dispersed activity when without larvae, which was true for both single n1* and grouped n6* ants (Wilcoxon signed rank exact test; n1 vs n1*: V=2, p-value=0.003, n6 vs n6*: V=5, p-value=0.01 (Figure 8-(c)). For single ants, we also found that its walking percentage was significantly higher when the ant was without larvae (and in a circular arena) than when it was with larvae (and in a square-shape arena). See Figure 8-(e), Wilcoxon signed rank exact test; n1 vs n1*: V = 0, p < 0.001. Here, walking was defined by the following criterion; the distance of two motion points in two consecutive frames (in one second) exceeded 120% of ant’s body length. Also, single ants without larvae walked longer distances (Figure 8-(f), Wilcox signed rank exact test; n1-mean vs n1*-mean: V=0, p < 0.001, n1-Median vs n1*-Median: V=1, p = 0.002, n1-Max. vs n1*-Max.: V=0, p < 0.001).

### 3.4 Response of groups to disturbance

By transferring the ants from their stock colony to the experimental filming arena, the ants experienced a sudden change in conditions. We therefore used our system to analyze whether their activity was different in the first period after the disturbance, and after which time activity levels would indicate an acclimation to the altered condition. We found that the ants, when in groups of 6 or 10, but not when alone, showed an initial high activity level, which then gradually subsided within 2-3 days (Figure 9-(a)). The regression lines in Figure 9-(a) show varying degrees of activity declines in most aggregates from different colonies. Figure 9-(b) shows a tendency of lower slopes with an increasing number of ants and in the presence of larvae, with significantly lower slopes occurring in groups of 6 and 10 ants compared to single ants (Kruskal-Wallis rank sum test, chi-squared = 22.337, df = 2, p <0.001; Dunn’s test of multiple comparisons using rank sums with Bonferroni correction: n1 vs n6, p = 0.001, n1 vs n10, p<0.001, n6 vs n10, p=0.963). The absence of larvae in n1* and n6* in the circular arena did not lead to a significant change in slopes compared to the single ant with larvae in the square-shaped arena (see Figure 9-(c); Wilcoxon signed rank tests; n1 vs n1*: V = 21, p = 0.32, n1 vs n6*: V = 53, p = 0.083). However, n6 groups with larvae in a square arena showed significantly lower slopes compared to n6 groups without larvae in a circular arena (Wilcoxon signed rank test, n6 vs n6*: V = 0 & p < 0.001). We further used ‘proxMCluster’ to analyze the bundled-up behaviour, measuring the activity intensity produced by multiple ants that are in close proximity (see Algorithm 1 in Appendix). Many ant groups showed distinctively high values at the very beginning, followed by smaller peaks throughout the week (Figure 10-(a)). However when larvae were absent, the ants’ overall activity when close to each other was significantly lower (Figure 10-(c)) and the otherwise observed distinctive high value at the very beginning was mostly absent (Figure 10-(b)). This significant lower activity within the proximate range to nestmates was observed while the overall activity was not significantly different (see section-3.1).

**Figure 9.**
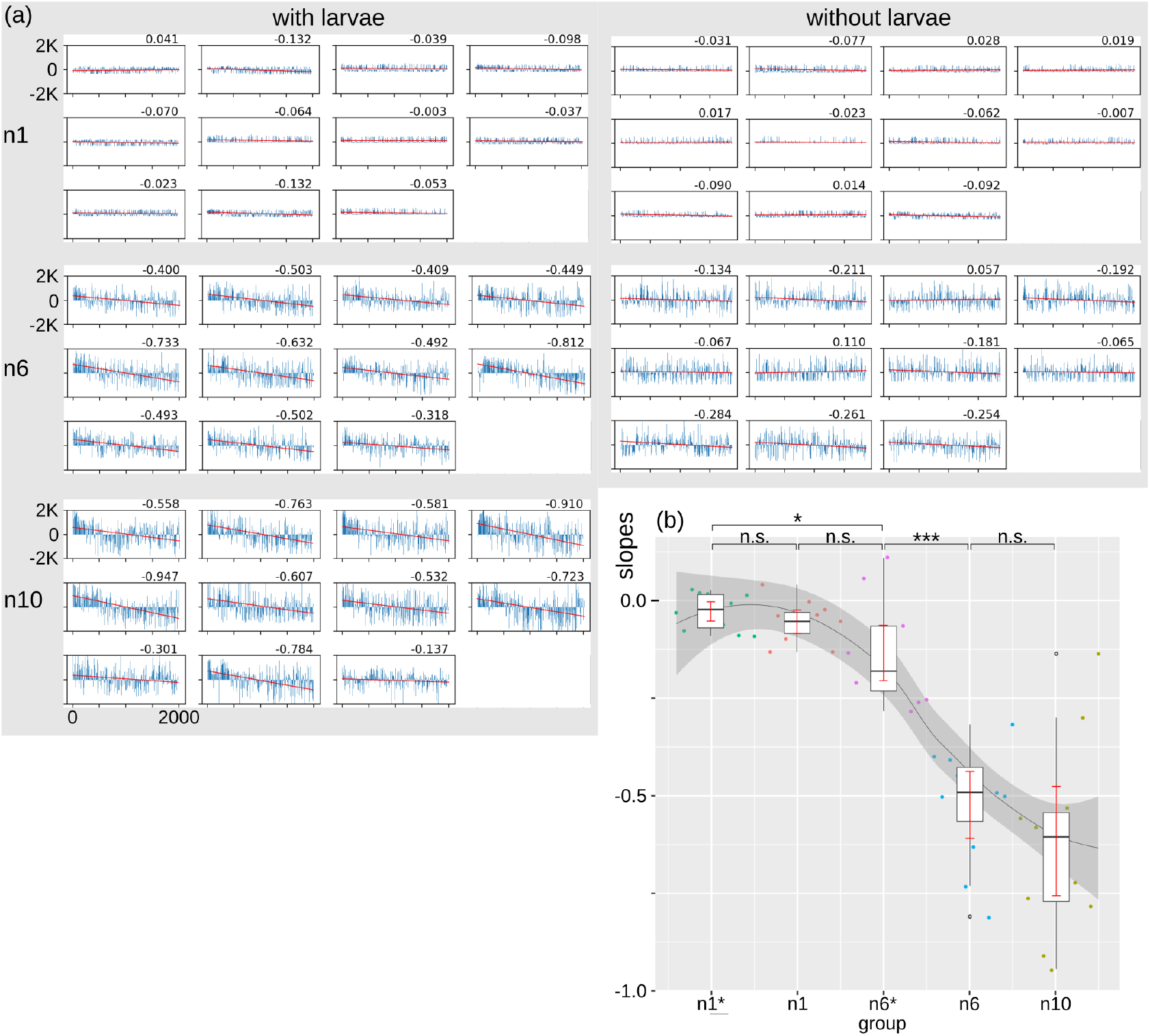
(a) Activity fluctuation for one week in single and grouped ants and in dependence of larval presence/absence. Zero on the y-axis represents the median of activity. Each vertical line shows the data for 5 minutes, and the red line shows the regression line over time, with the number indicating its slope. (b) Comparisons of the regression line slopes across group sizes and larval presence. Boxplots are formatted as in Figure 7, except dots. Dots here are data points differently coloured for each group.

**Figure 10.**
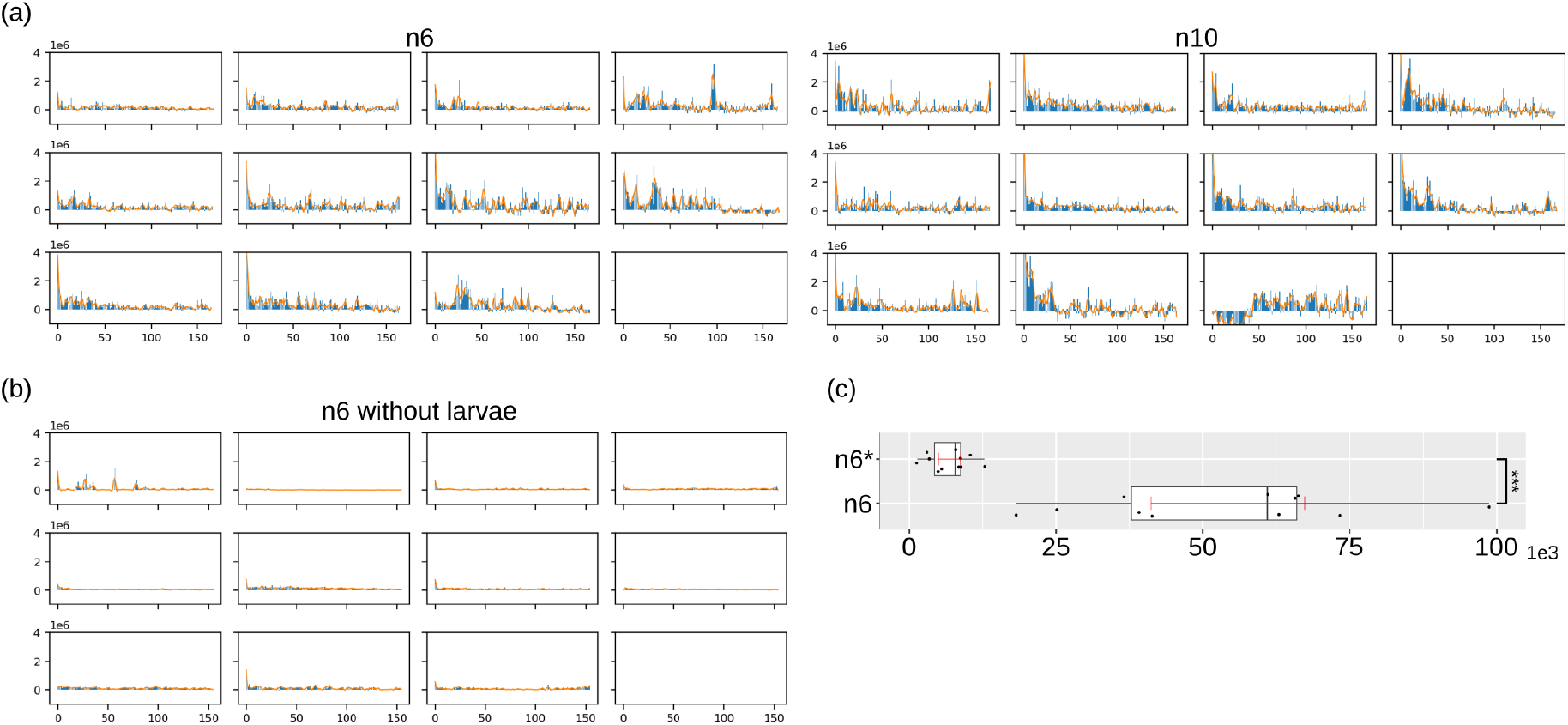
Bundled activity intensity. Graphs of ‘proxMCluster’ with zero on the y-axis representing the median. An individual vertical line shows the data for one hour. (a) Graphs of groups of 6 and 10 ants with larvae. (b) Graph of the group of 6 ants without larvae. (c) Comparison of mean values of n6 and n6* (Wilcoxon signed rank exact test; V = 66, p-value < 0.001). Boxplots are formatted as in Figure 7.

### 3.5 Simulation results

We used AntSim to simulate the activity of individual ants and examine how simple rules at the individual level could reproduce group-level patterns observed in our empirical data. Specifically, we aimed to capture two key features: (1) temporal fluctuations in activity (3.2) and (2) reduced overall activity levels in the n10 group (3.1).

To ensure that our simulations reflected steady-state behavior rather than the initial transitional effect, we restricted the simulation result comparison to a 96-hour period following an initial 60-hour acclimation phase, based on observations of real ants adjusting to the flat-nest environment. We began by extracting activity and inactivity duration distributions from single-ant data (n1, *n* = 11), computing the average probability distribution across individuals. The simulations based solely on these average distributions failed to reproduce the patterns observed in real data.

This discrepancy arose because averaging across individuals led to an underrepresentation of longer activity and inactivity durations, which are essential for capturing the variability and stochasticity observed in real ants. To address this, we applied a probabilistic “boosting” to the tail of the duration distributions for each simulated ant. Specifically, for each individual, we selected a threshold duration by adding a small random deviation to a group-level reference value, which was itself randomly chosen within a predefined range (0*–* 300*s* for activity, 600*–* 1200*s* for inactivity, based on empirical observations). We then increased the probability mass of durations exceeding this threshold by applying a multiplicative scaling factor, iteratively adjusted until the total tail probability reached at least 0.05. This adjustment allowed the simulations to retain the overall statistical structure of the group-level data while introducing individual variability necessary to reproduce the long-tailed distributions observed in real ants. Adding this boosting step improved the fit between simulated and real ants’ activity patterns (Figure 11-(a)). Statistical comparison using the Wilcoxon rank-sum test showed that simulation without boosting (sim.1) differed significantly from real data (W = 23, p < 0.001), while simulations with boosting but no interaction (sim.2) showed no significant difference (W = 432, p = 0.152).

**Figure 11.**
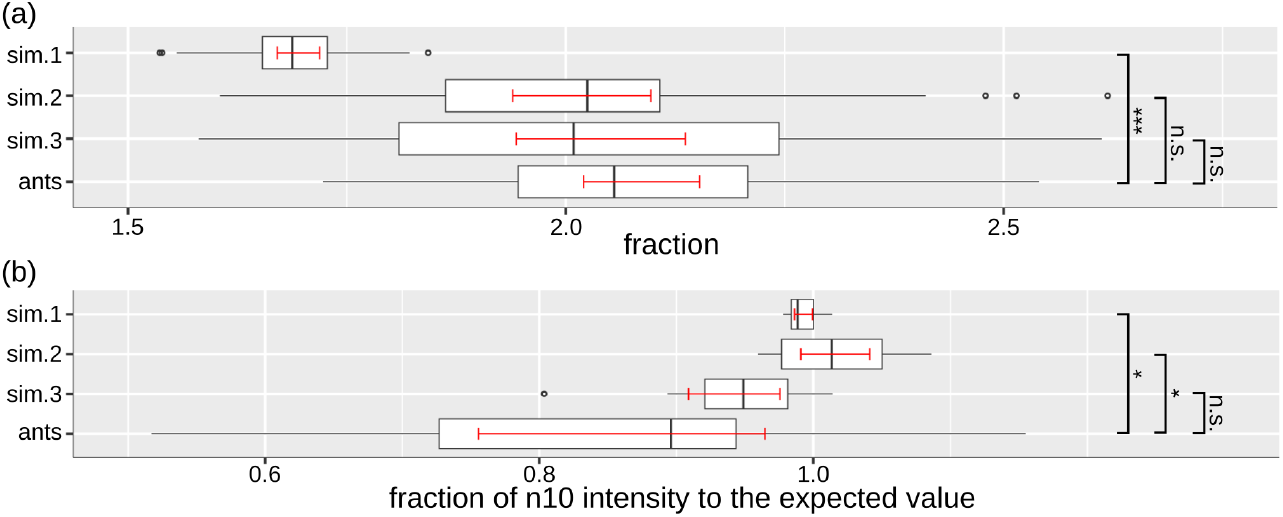
Comparison of simulated data to the real-world data; sim.1: No boosting & No interaction, sim.2: Boosting & No interaction, sim.3: Boosting & Interaction, ants: Real-world data. (a) Comparison of fractions in the longer to the shorter period range. (b) Comparison of fractions of n10 activity to the expected value by linear increase from n6. Boxplots are formatted as in Figure 7.

However, sim.2 still failed to replicate the reduced activity levels observed in n10. To address this, we introduced an interaction rule: when many ants in the group were active simultaneously, the probability of longer inactivity durations increased for each individual. This rule was motivated by the hypothesis that ants may reduce their own activity when others are already active, possibly reflecting redundancy or energy conservation mechanisms. With this addition (sim.3: boosting + interaction), the simulation reproduced both the activity fluctuation pattern and the reduced activity intensity in the n10 group. The Wilcoxon rank sum tests showed no significant differences between real data and sim.3 for both overall activity fluctuation (W = 454, p = 0.25) and n10 inactivity patterns (Figure 11-(b); W = 83, p = 0.151).

In summary, our simulation approach was built bottom-up, starting from simplified individual-level rules and progressively incorporating minimum essential features to reproduce the collective behavior patterns observed in real ants.

## 4 Discussion

We have developed a multicomponent system that can be utilized for studying long-term behavioural changes of groups of ants or other insects. It is a collection of tools, consisting of (i) a custom-designed nest preventing occlusion during data acquisition, (ii) a recording system running on a single-board-computer and (iii) several programs in Python for data acquisition, analysis and simulation. All software and design (Blender) files of the flat-nest are available via the Github repository https://github.com/jinook0707/CremerGroupApp.AntOS, Visualizer and AntSim are located in ‘antOS’, ‘visualizer’ and ‘antSim’ subfolders respectively. The design files are located in ‘blenderFiles’ folder. We here showcase the use of our system by an experimental study comparing the activity of *L. niger* worker ants differing in group size (1, 6, 10 ants), presence/absence of larvae and nest architecture (square vs circular). The use of AntOS allowed us to derive the following behavioural characteristics from our experiment:

1. The activity intensity (amount of motions) in ant groups was found to be lower than the expected value of the linear increase with the number of workers.

2. A weak periodicity in the lower frequency range (longer duration range; 20-120 minutes) was observed independent of group size. Because this weak periodicity did not show a particular peak on a specific duration, we used a fraction of the long duration range to the short duration range, which was 2.032, 2.041 and 2.102 (median values) in n1, n6 and n10, respectively. In the absence of larvae (in a circular arena), the periodicity became even weaker and the fraction dropped to 1.552 and 1.781 for both single (n1*) and groups of 6 ants (n6*). This suggests that a weak periodicity occurs in *L. niger* workers, which is promoted further by presence of larvae or nestmates, with the latter having a less strong effect than brood presence.

3. Workers are spatially attracted to larvae, edges (or walls), the water-source and nestmates.

4. Single workers with and without larvae showed similar activity levels (total amount of motions). Yet, in the presence of larvae, the ant seems to spend more time for non-walking behaviours such as grooming and care-taking. In the absence of larvae, it spends more time walking, similar to the report from other ant species [49] in which isolated ants without any other individual moved longer distances.

5. Ant groups, but not isolated individuals displayed higher activity in the beginning of the experiment, and decreased their activity levels over 2-3 days to reach a stable activity level after acclimation to the experimental condition. Notably, this increased initial activity was dependent on larval presence. Although the number of workers in the group might affect it to certain degree, it was statistically not significant. Larval presence also triggered higher activity of workers in close proximity to their nestmates. This suggests that ant groups with larvae in a new environment first highly increase their activity, keeping short distances to other nestmates, then gradually increasing the distance between nestmates, likely for exploration of the new space, and gradually decreasing the activity level to a stable state.

Our experimental findings were further supported by generating data by simulation based on several theoretical rules inferred form the real-world ant data using Visualizer. Implementation of the rules in AntSim and verification, which simulation rules are required to produce results that match the observed experimental data, allows further interpretation of the results and its underlying biological rules. Currently, AntSim only produces reliable data for temporal but not yet spatial activity patterns, which will require further experiments and methodological developments.

Our system can also be extended by adding further sensors and actuators to the SBC to enable different types of automatic data collection and experimental manipulation such as e.g. automatic increase/decrease of temperature and humidity.

Whilst we developed this method to be independent of individual tracking, AntOS includes a built-in colour detection feature that can be used to record the positions of worker groups and a focal individual ant marked with the established colour-marking techniques [50]. This functionality could be extended to track multiple individuals within a colony by combining colour detection with additional algorithms such as blob segmentation, k-means algorithm, trained classifier, etc. The number of individuals that can be tracked, will depend on their body size and hence the number of color dots that can be applied to their body. In the case of *L. niger*, likely only two colour dots of large enough size can be placed on each ant, which given with the limited range of colours that can be differentially recorded in the system (approx. 4 *–* 5) limits the number of traceable colour coded individuals to approx. 20 in our system.

As showcased in our experimental study, AntOS already provides a powerful tool to observe and analyze long-term temporal and spatial collective activity, including the emergence of subtle changes and reaction to disturbance over time. As such, we consider the framework developed in this study to be a versatile analysis tool allowing in-depth analysis of otherwise hard-to-grasp noisy data of behavioural patterns of social insect colonies or other insect groups as a whole, and to pick up gradual changes over a long-term period. Importantly, it does not require tagging of individuals nor does it rely on the expression of very clear behavioural changes. Due to its simplicity of hardware building blocks and low requirements, the system is easily scalable as it allows parallel, continuous observation of a high number of colonies. It can therefore be built up as a high-throughput system to simultaneously analyze high replicate numbers of colonies over long periods for both their temporal and spatial collective activity patterns. Our system can be used to follow the colony development and growth as a direct estimate for fitness, and how it is influenced by different experimental conditions.

## Data availability statement

All code and data are provided under Github https://github.com/jinook0707/CremerGroupApp.

## Author contributions

Jinook Oh and Sylvia Cremer conceived the project together. Jinook OH designed and built the hardware and software, collected and analysed the data. Jinook Oh led the writing of the manuscript, Sylvia Cremer contributed critically to the drafts and both authors gave final approval for publication.

## Acknowledgements

We thank Harikrishnan Rajendran for discussion and the European Research Council (ERC) under the European Union’s Horizon 2020 Research and Innovation Programme for funding (No. 771402; EPIDEMICSonCHIP to SC).

**Appendix**

## APPENDIX A Python files for software of the framework

**Figure A1.**
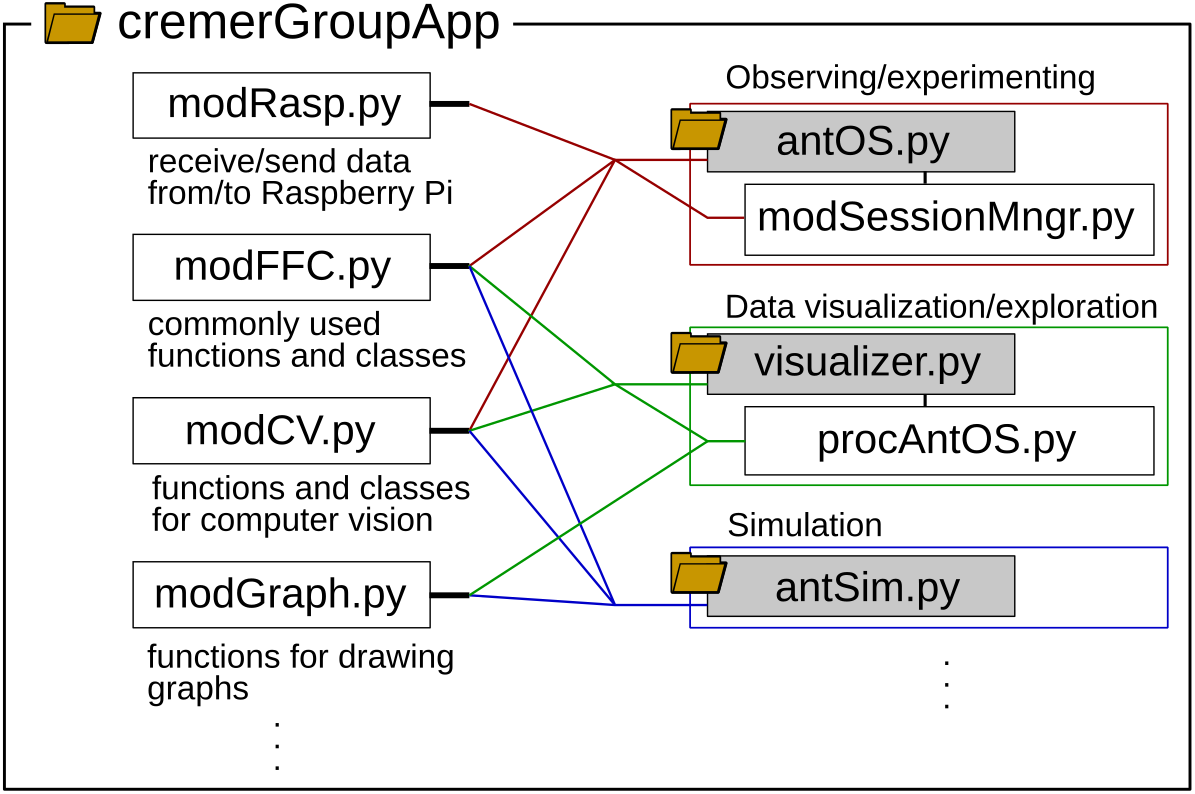
Python files for the three programs used in this study

## APPENDIX B Commonly used classes & functions

The following tables are to list major functionalities implemented in shared script files.

**Table B1.** modFFC.py.

| Functionality | Function (class) names |
| --- | --- |
| GUI processing | set_img_for_btn, getWXFonts, setupPanel, addWxWidgets, updateFrameSize, calcImgSzFitToPSz, add2gbs, widgetValue, preProcUIEvt, showStatusBarMsg, procExceptionWX, calcI2DIRatio |
| Image & sound | getConspicuousCol, wxSndPlay |
| Thread run in wx-Python | startTaskThread, postProcTaskThread, makeWaitingMessagePanel, receiveDataFromQueue |
| Python logging | setMyLogger |
| File/folder path | chkFPath, getFilePaths_recur |
| Time-stamp | get_time_stamp, get_datetime |
| Save string to a file | writeFile |
| Processing number & geometry | str2num, convt_idx_to_ordinal, convt_180_to_360, convt_360_to_180, calc_pt_w_angle_n_dist, calc_line_angle, calc_angle_diff, calc_pt_dists, calc_pt_line_dist, rot_pt, convert-SagittaArc2AngArc |
| NumPy array processing | csv2numpyArr, getNumpyDataType |
| Outlier detection | OutlierDetection_GESD (class) |
| Custom dialog | PopupDialog (class) |

**Table B2.**
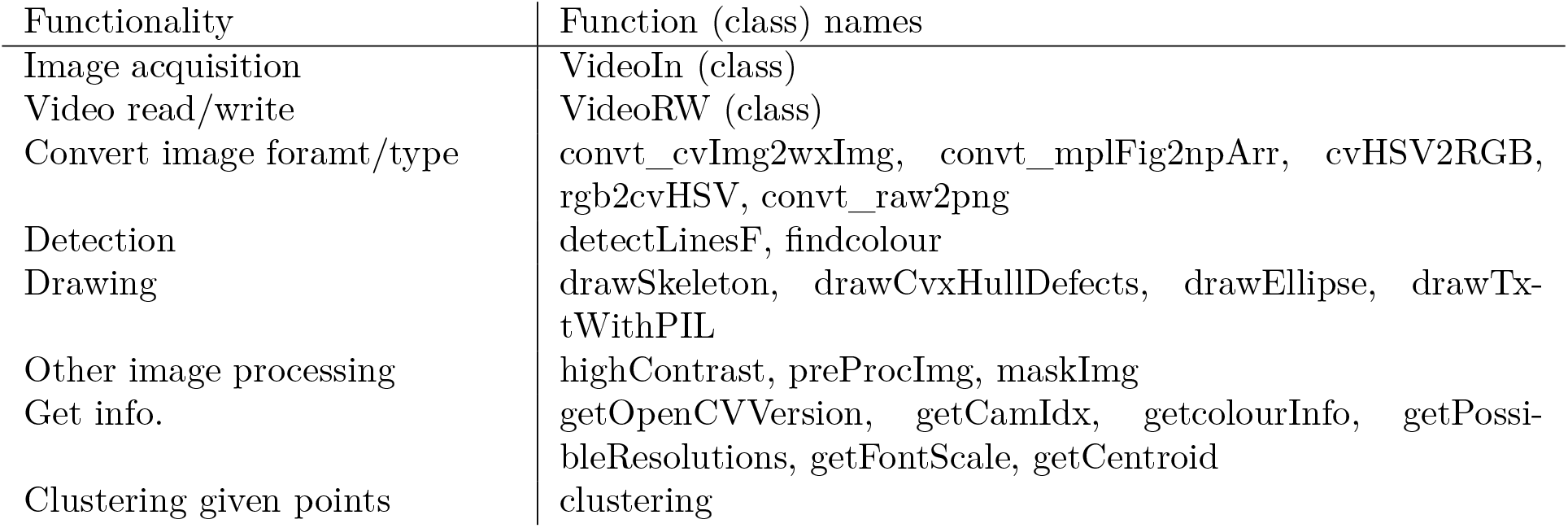
modCV.py.

| Functionality | Function (class) names |
| --- | --- |
| Image acquisition | VideoIn (class) |
| Video read/write | VideoRW (class) |
| Convert image format/type | convt_cvImg2wxImg, convt_mplFig2npArr, cvHSV2RGB, rgb2cvHSV, convt_raw2png |
| Detection | detectLinesF, findcolour |
| Drawing | drawSkeleton, drawCvxHullDefects, drawEllipse, drawTxtWithPIL |
| Other image processing | highContrast, preProcImg, maskImg |
| Get info. | getOpenCVVersion, getCamIdx, getcolourInfo, getPossibleResolutions, getFontScale, getCentroid |
| Clustering given points | clustering |

**Table B3.** modRasp.py.

| Functionality | Function (class) names |
| --- | --- |
| Processing with Raspberry Pi GPIO pins | RaspGPIO (class) |
| Temperature logging | RaspGPIO.logTemp |
| Send a signal to a pin | RaspGPIO.sendSignal |
| Image acquisition on Raspberry Pi CSI camera | RaspCSICam (class) |

**Table B4.**
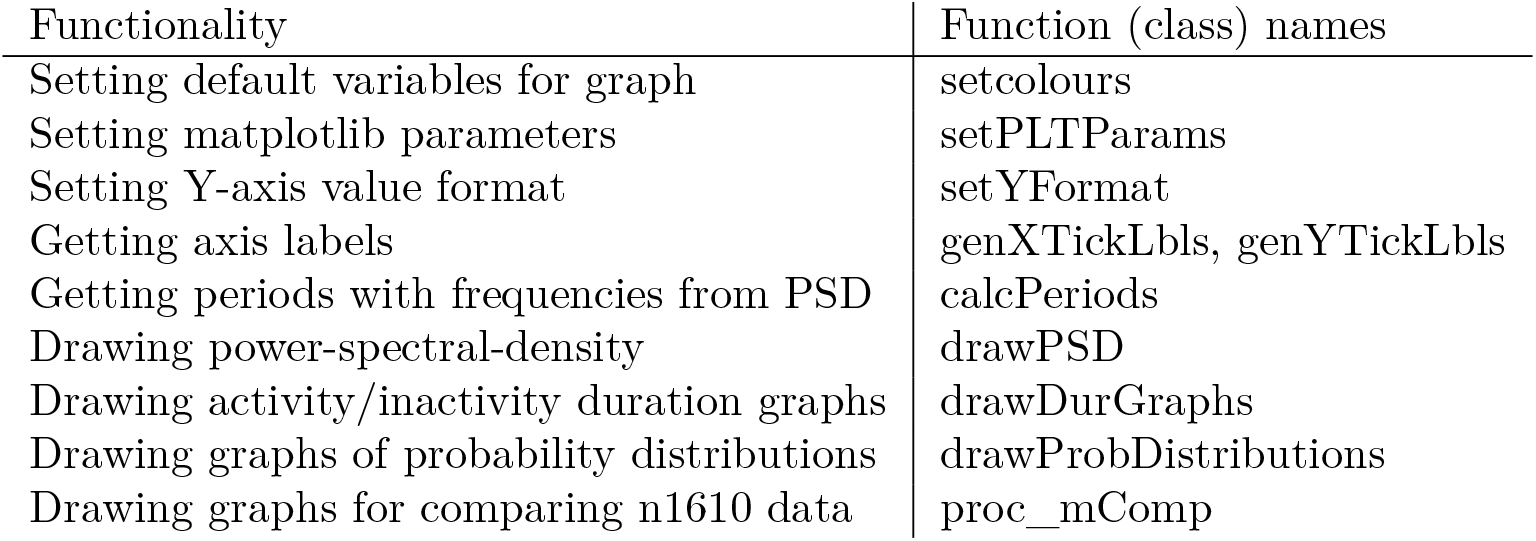
modGraph.py.

## APPENDIX C Calculating ’proxMCluster’ values

### Algorithm 1

*proxMCluster*’

^*^A minimum duration (occurring in two consecutive bundled data) was set to rule out accidental co-occurrences of close motions.

^*^ ’maxNOfOneAnt’ was set to five. Each motion point is a center point of a motion contour which does not necessarily mean an entire body of an ant. Multiple motion points can be produced by a single ant, such as one point on its antenna, a second a point on a leg, etc. When analyzing single ants each for one week (n = 11 replicates), 99.6 % of the motion points per frame were in the range of 1-5 (0.394, 0.358, 0.169, 0.06, and 0.015 for 1-5 motion-points respectively).

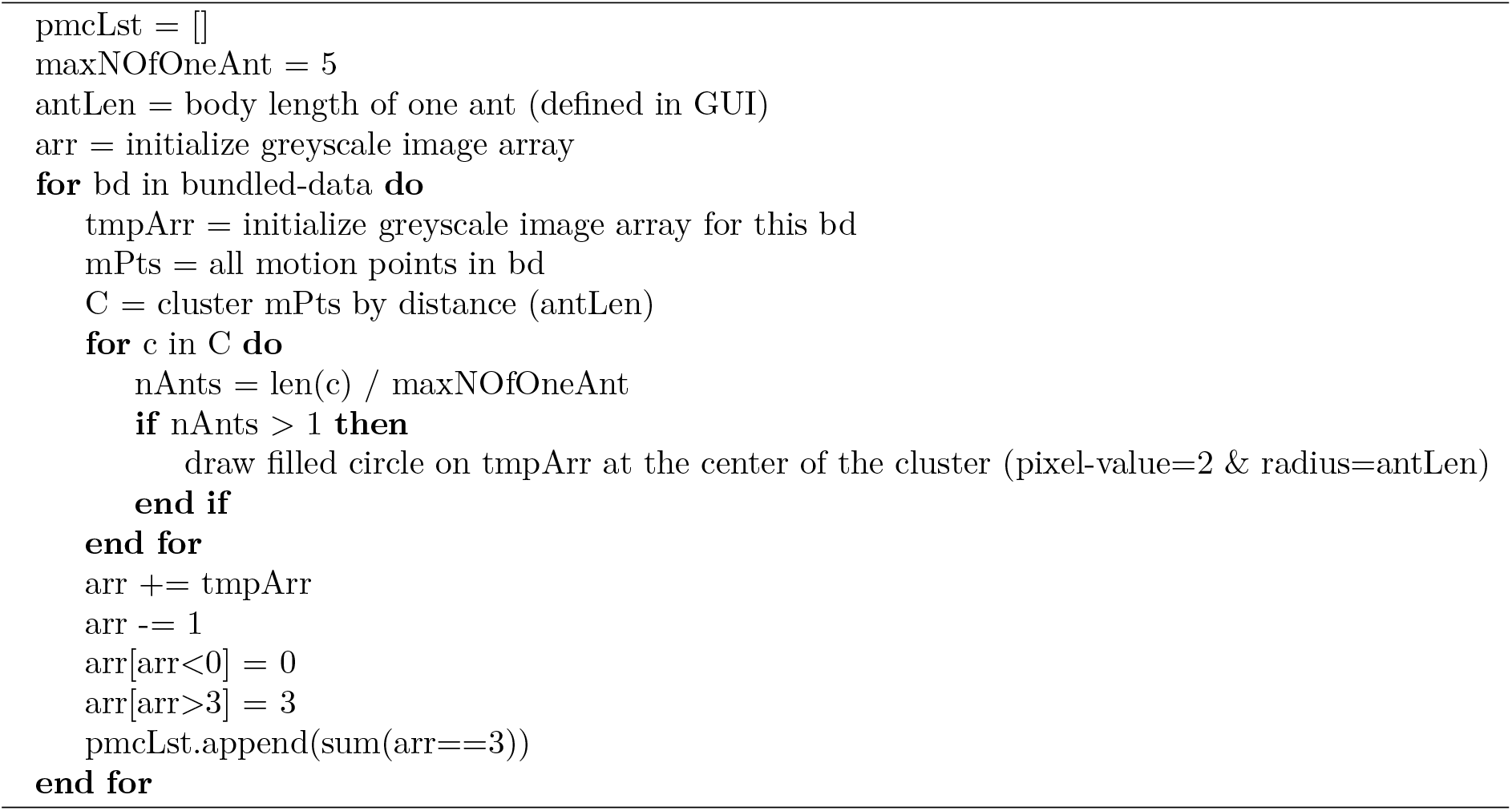

